# Cell viability is dominated by quantum effects

**DOI:** 10.1101/2023.07.18.549596

**Authors:** Takeshi Yasuda, Nakako Nakajima, Tomoko Yanaka, Takaya Gotoh, Wataru Kagawa, Kaoru Sugasawa, Katsushi Tajima

## Abstract

Quantum tunneling is a phenomenon in which small quantum particles pass through a reaction energy barrier, as if they were passing through a tunnel opened in the barrier. In this study, we analyzed the involvement of the quantum tunneling effect in enzymatic chemical reactions involving hydrolysis *in vitro*, by monitoring the kinetic isotope effects due to hydrogen isotopes and their temperature dependence as indicators. The results demonstrated that the quantum tunneling effect is involved in deacetylation, DNA cleavage, and protein cleavage reactions. These related reactions were also examined in terms of their effects on cells, which revealed that the quantum effect is even involved in cell survival, including the almost complete inhibition of DNA homologous recombinational repair.

**One-Sentence Summary:** Due to quantum effects in numerous enzymatic reactions mediated by hydrolysis, heavy water significantly impacts biological outcomes in cells.

## Introduction

Quantum biology is an exploratory science that seeks to understand biological life phenomena from the standpoint of quantum mechanics theory. The concept is described in the excellent book “Solving the Mysteries of Life with Quantum Mechanics” by Jim Al-Khalili and Johnjoe McFadden (*1*). A quantum is a very small discrete unit of matter or energy, such as a proton, electron, or photon, with both particle and wave properties. Therefore, in quantum biology, the biological processes mediated by photons, such as photosynthesis, photorepair, and photoreceptor signaling, are examined and interpreted by quantum theory (*2-4*).

A variety of proteins catalyze chemical reactions with DNA and protein substrates. In the enzyme-catalyzed cleavage or formation of a chemical bond in the substrate, the electron involved in the chemical bond is expected to behave quantum mechanically. The starting reactant exists in the lowest energy state, called the ground state (GS). For the chemical reaction to proceed, it must overcome an activation energy barrier, in which the highest energy state is the transition state (TS). The activation energy barrier can be surmounted by heat in a chemical reaction. A catalyst promotes a chemical reaction without heating, by forming a reaction intermediate complexed with the starting material, and thereby provides a new reaction pathway with a lower activation energy barrier. In quantum phenomena, quantum particles can pass through the barrier. Therefore, the chemical reaction proceeds without reaching the transition state. This phenomenon is called “quantum tunneling”. The effect of quantum tunneling can be modified by substituting the atom directory involved in the chemical reaction with its isotope. More energy is required to cleave the chemical bond involving the heavier isotope; therefore, the reaction rate is decreased by the substitution with the heavier isotope (*5-6*). This phenomenon is called the kinetic isotope effect. Previously, by using the kinetic isotope effect as a probe, the quantum tunneling effect was examined in proteins, such as dehydrogenases (*7-8*), which catalyze a hydrogen-transfer reaction during substrate oxidation. In this reaction, the kinetic isotope effect was examined by comparing the transfer of normal hydrogen (^1^H or H) and its heavier isotope deuterium (^2^H or D) (*8*). The transfer of normal hydrogen by the enzymes was faster than that of deuterium, indicating the quantum tunneling effect in enzyme reactions.

Although the quantum tunneling effect in dehydrogenases has been reported, little is known about its involvement in other types of enzymes with various functions. In addition, the biological effects derived from the quantum tunneling have not been clarified. Therefore, this study aimed to elucidate the quantum effects of enzymes related to various important biological phenomena and their associated effects on cells.

## Results

### Concept of the research

The chemical reaction of the reactant mediated by quantum particles at the GS can proceed without reaching the TS by the quantum tunneling effect (fig. S1A). Because deuterium (D) has a lower probability of quantum tunneling than hydrogen (H), a kinetic isotope effect appears in enzymatic chemical reactions involving hydrogen, in which replacing H with D results in a lower reaction rate (fig. S1B, text S1). The connection between quantum tunneling and kinetic isotope effects can be understood from the equation for the probability of the quantum tunneling effect, which is derived from the Schrödinger equation (text S1) described in textbooks, such as that by P.W. Atkins (*9*). The probability of the quantum tunneling effect of D is lower than that of H, thereby resulting in the kinetic isotope effects. The kinetic isotope effect is also caused by differences in the vibrational potential energies of the chemical bonds involved in the reaction, which are also related to quantum mechanics (fig. S1C, text S2) (*5-6, 9*). When the reaction temperature is high, the reaction can proceed beyond the potential energy peak without quantum tunneling. Therefore, the quantum tunneling effect is reduced or eliminated at higher temperatures, and accordingly, this temperature dependency is also an indicator (*5-6, 9*).

### Kinetic isotope effect on SIRT3-mediated human RAD52 deacetylation

Hydrogen-transfer is involved in deacetylation (*10-12*), which is the reverse reaction of acetylation. During deacetylation, a hydrogen of a water molecule directly attacks the chemical bond of acetyl-lysine to remove the acetyl group, and is transferred to the deacetylated lysine (fig. S1D). Acetylation is catalyzed by histone acetyltransferases (HATs), and deacetylation is accomplished by histone deacetylases (HDACs). Human RAD52, a DNA double strand break (DSB) repair protein (*13*), is acetylated by a HAT, p300, and is deacetylated by an HDAC, SIRT3 (*14*). To examine whether the deacetylation reaction is subjected to the kinetic isotope effect, we performed *in vitro* RAD52 deacetylation by SIRT3 (*14*). At first, RAD52 is incubated at 30°C with p300 and Acetyl Coenzyme A (Ac-CoA) in reaction buffer containing H_2_O or D_2_O, for the RAD52 acetylation (HAT assay). Because RAD52 acetylation is inhibited in the presence of single stranded (ss)DNA (*14*), ssDNA is then added to stop the ongoing RAD52 acetylation reaction. A portion of the reaction mixture containing the acetylated RAD52 is incubated with SIRT3 and NAD^+^ in buffer containing H_2_O or D_2_O, at 30°C (HDAC assay) (fig. 1A). Each sample before and after the HDAC assay was examined by immunoblotting (fig. 1B). The band intensities of Ac-RAD52 before the HDAC assay were almost the same between the samples incubated in the presence of H_2_O and D_2_O (fig. 1, B and C). In contrast, after the HDAC assay, the relative intensity of the Ac-RAD52 band was more efficiently decreased in the presence of H_2_O than in the presence of D_2_O (fig. 1, B and C). Therefore, the presence of D_2_O decreased the SIRT3-mediated deacetylation reaction of Ac-RAD52.

**Fig. 1.**
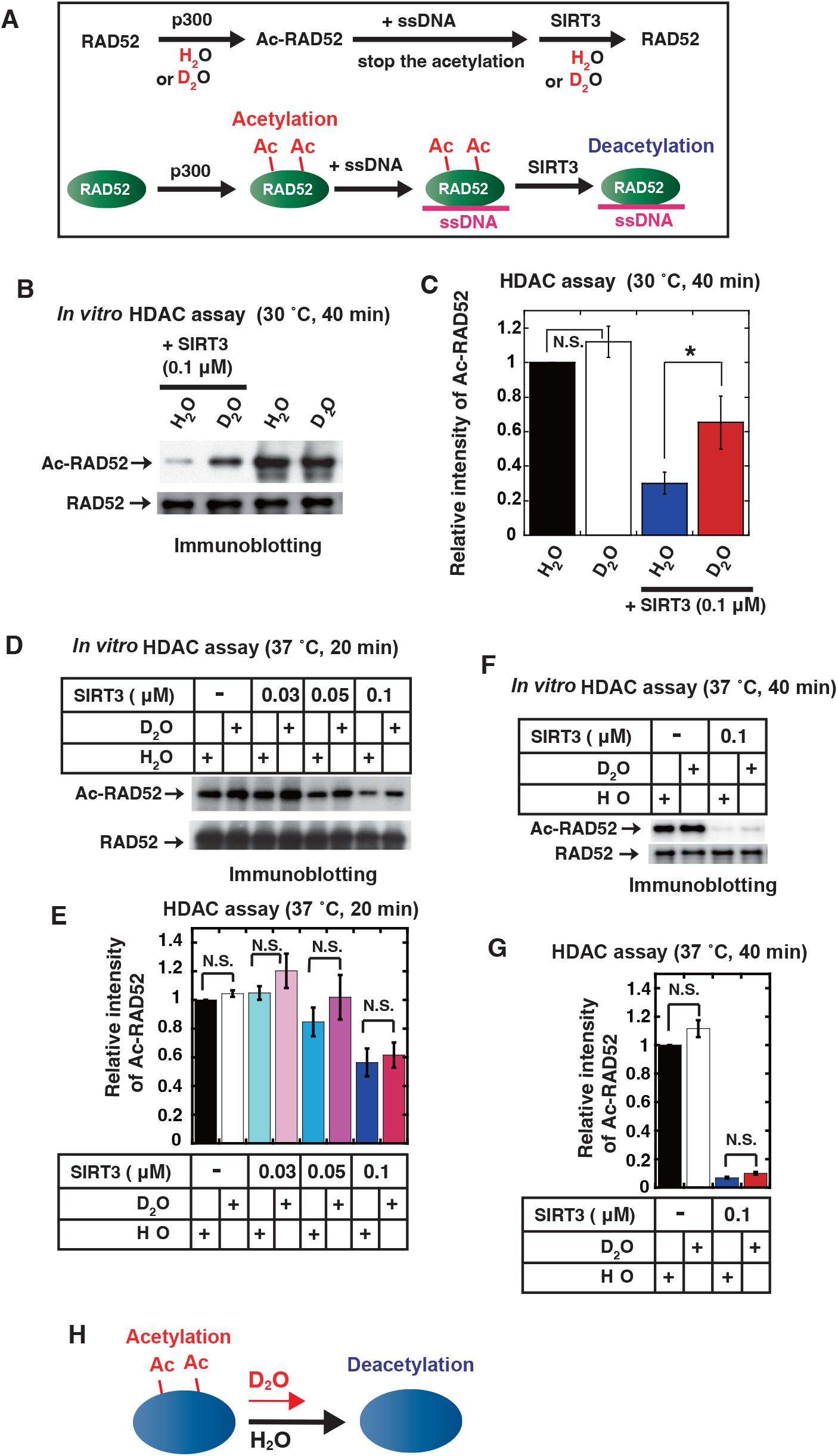
Kinetic isotope effect on RAD52 protein deacetylation by SIRT3. (**A**) Schematic representation of the RAD52 deacetylation assay. (**B, D**, and **F**) *In vitro* acetylation assays of RAD52 were performed, as described in the Materials and Methods. After the addition of a poly dT 68 mer, aliquots of the reaction mixture containing RAD52 proteins (0.1 μM in Fig. 1B and 0.08 μM in Fig. 1D and Fig. 1F) were incubated with the indicated amount of SIRT3 in HDAC buffer in the presence of H_2_O or D_2_O, at the indicated temperature for the indicated times. The reaction mixtures were subjected to SDS-PAGE, followed by immunoblotting with an anti-acetylated lysine antibody (top: Ac-RAD52) and an anti-RAD52 antibody (bottom: RAD52). (**C, E**, and **G**) Relative band intensities of acetylated RAD52 normalized to those of the RAD52 bands. The graph shows the mean values and standard errors of the mean from 4 (C) or 3 (E and G) independent experiments. The samples connected by lines were compared (\**P* <0.05 and N.S., not significant by an unpaired Student’s t-test). (**H**) Schematic representation of the experimental results for the kinetic isotope effect on the protein deacetylation reaction.

Next, we conducted the same experiments using H_2_O and D_2_O obtained from several different manufacturers (fig. S2, A and B). As a result, there were no significant differences in the deacetylation rates among the three types of H_2_O-containing buffers or between the two types of D_2_O-containing buffers. Therefore, regardless of the types of H_2_O or D_2_O, the effect on the HDAC reaction rate was solely caused by the difference in the isotope between H_2_O and D_2_O. When the SIRT3-mediated RAD52 deacetylation assays were performed at 42°C, the kinetic isotope effects disappeared (fig. 1, D to G). This temperature dependency suggests the involvement of the quantum tunneling effect in the SIRT3-mediated RAD52 deacetylation (fig. 1H).

### Kinetic isotope effect on proteolytic protein cleavage

Protein cleavage by protease family proteins is an enzymatic reaction mediated by hydrolysis. The protease family proteins are involved in diverse biological functions, such as protein degradation, signal transduction, apoptosis, autophagy, immunity, and viral infection (*15-19*). We examined the involvement of the quantum effect on protein cleavage by caspase-3, a protease that functions in apoptosis induction (*18*) (fig. 2, A-E). The caspase-3 substrate is the DEVD peptide sequence, and its derivative, Ac-DEVD-AMC, generates fluorescence upon cleavage (*18, 20*) (fig. 2D). The time-course of DEVD cleavage was monitored with H_2_O or D_2_O in the reaction buffer, at 23°C or 37°C (fig. 2, A and B). At both reaction temperatures, the reaction rate was lower with D_2_O than with H_2_O (fig. 2, A, B and E). The slope ratio in the presence of H_2_O to D_2_O was 1.75 at 23°C, and decreased to 1.48 at 37°C (fig. 2, A and B). Therefore, the temperature dependency of the difference in the kinetic isotope effects suggests the involvement of the quantum tunneling effect in the caspase-3-mediated peptide cleavage reaction.

**Fig. 2.**
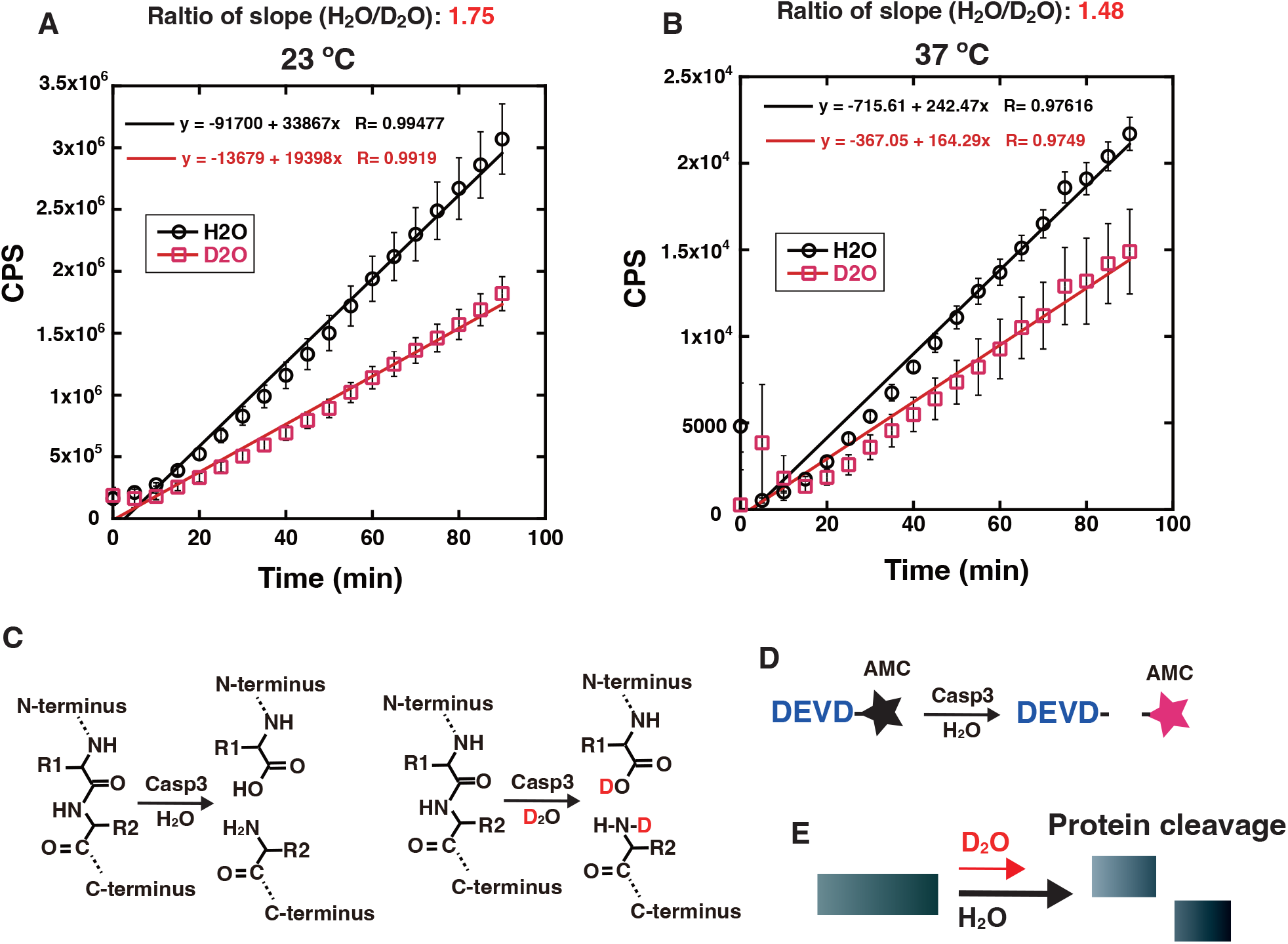
Kinetic isotope effect and its temperature dependency in peptide-bond cleavage by caspase-3. (**A** and **B**) *In vitro* caspase-3-mediated peptide-bond cleavage assays were performed at 23°C (A) or 37°C (B) in the presence of H_2_O or D_2_O, as described in the Materials and Methods. The graph shows the mean values and standard errors of the mean from 3 samples, with linear curve fitting. The ratios of the slopes of the linear curve fitting of H_2_O to D_2_O are shown at 23°C (A) and 37°C (B). (**C**) Schematic representation of peptide bond hydrolysis in the presence of H_2_O or D_2_O. (**D**) Schematic representation of the caspase-3-mediated peptide cleavage assay using the Ac-DEVD-AMC substrate. (**E**) Schematic representation of the experimental results for the kinetic isotope effect on protein cleavage.

### Kinetic isotope effect on nuclease-mediated DNA cleavage

DNA cleavage by nuclease family proteins is also mediated by hydrolysis (fig. 3A). The nuclease family proteins play a variety of roles, including DNA repair and recombination (*21-25*). We examined the involvement of the quantum effect on DNA cleavage by a nuclease, I-SceI (*26*). The cleavage rates by I-SceI against a linearized double-stranded DNA containing the specific cleavage sequence were examined in reaction solutions containing H_2_O or D_2_O (fig. 3B-D). Kinetic isotope effects were detected at 15°C, but disappeared at 37°C (fig. 3B-E). Thus, the kinetic isotope effect and the temperature dependency also suggest the involvement of the quantum tunneling effect in the I-SceI-mediated DNA cleavage reaction.

**Fig. 3.**
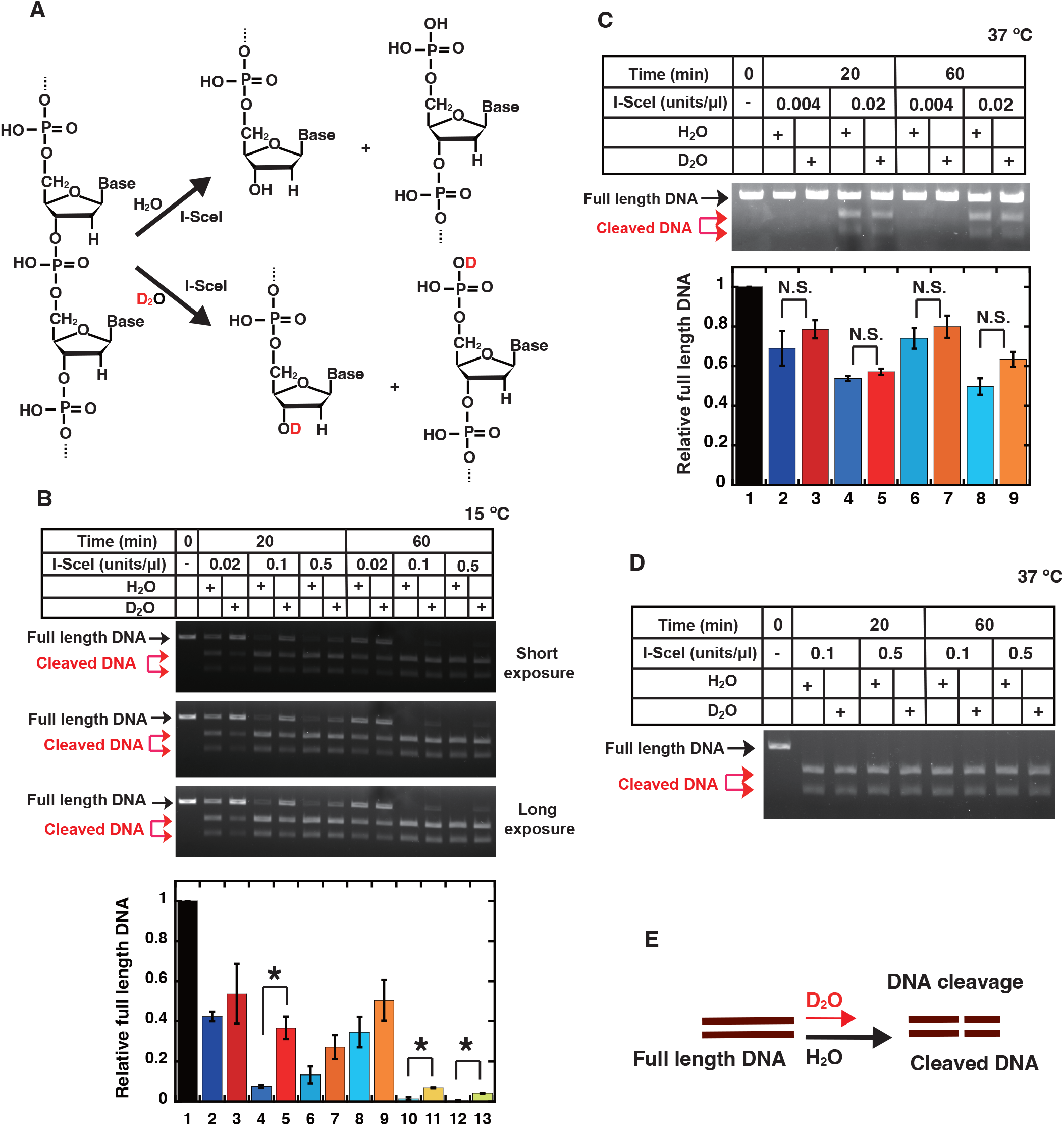
Kinetic isotope effect and its temperature dependency in DNA cleavage by I-SceI nuclease. (**A**) Schematic representation of phosphodiester bond hydrolysis by I-SceI in the presence of H_2_O or D_2_O. (**B** to **D**) *In vitro* I-SceI-mediated DNA cleavage assays were performed at 15°C (B) or 37°C (C and D) for the indicated times in the presence of H_2_O or D_2_O, as described in the Materials and Methods. (**B** and **D**) The relative band intensities of full-length DNA are shown in the graph. The mean values and standard errors of the mean from 3 independent experiments were plotted. The samples connected by lines were compared (\**P* <0.05 and N.S., not significant by an unpaired Student’s t-test). (**E**) Schematic representation of the experimental results for the kinetic isotope effect on DNA cleavage.

### Quantum effects severely impact HR repair required for protection of cell survival

To clarify the biological consequences of the quantum effects, we examined the isotope effects in cells with respect to the kinetic isotope effects of the SIRT3 and I-SceI enzymes detected in the *in vitro* experiments. For this purpose, we performed the HR repair assay using human DR-GFP reporter cells (fig. 4, A and B) (*27-28*). The I-SceI sequence is absent in the genome of normal human cells, and the DR-GFP reporter cells contain a reporter DNA cassette containing the I-SceI sequence in the genome. In the HR repair assay, a DSB is specifically induced within the reporter cassette when the I-SceI enzyme is produced by the expression plasmid transfected in the cells, and normal GFP genes are expressed when the DSB site is repaired by HR (fig. 4B). The HR repair frequency is dependent on the DSB induction frequency by I-SceI (fig. 4C).

**Fig. 4.**
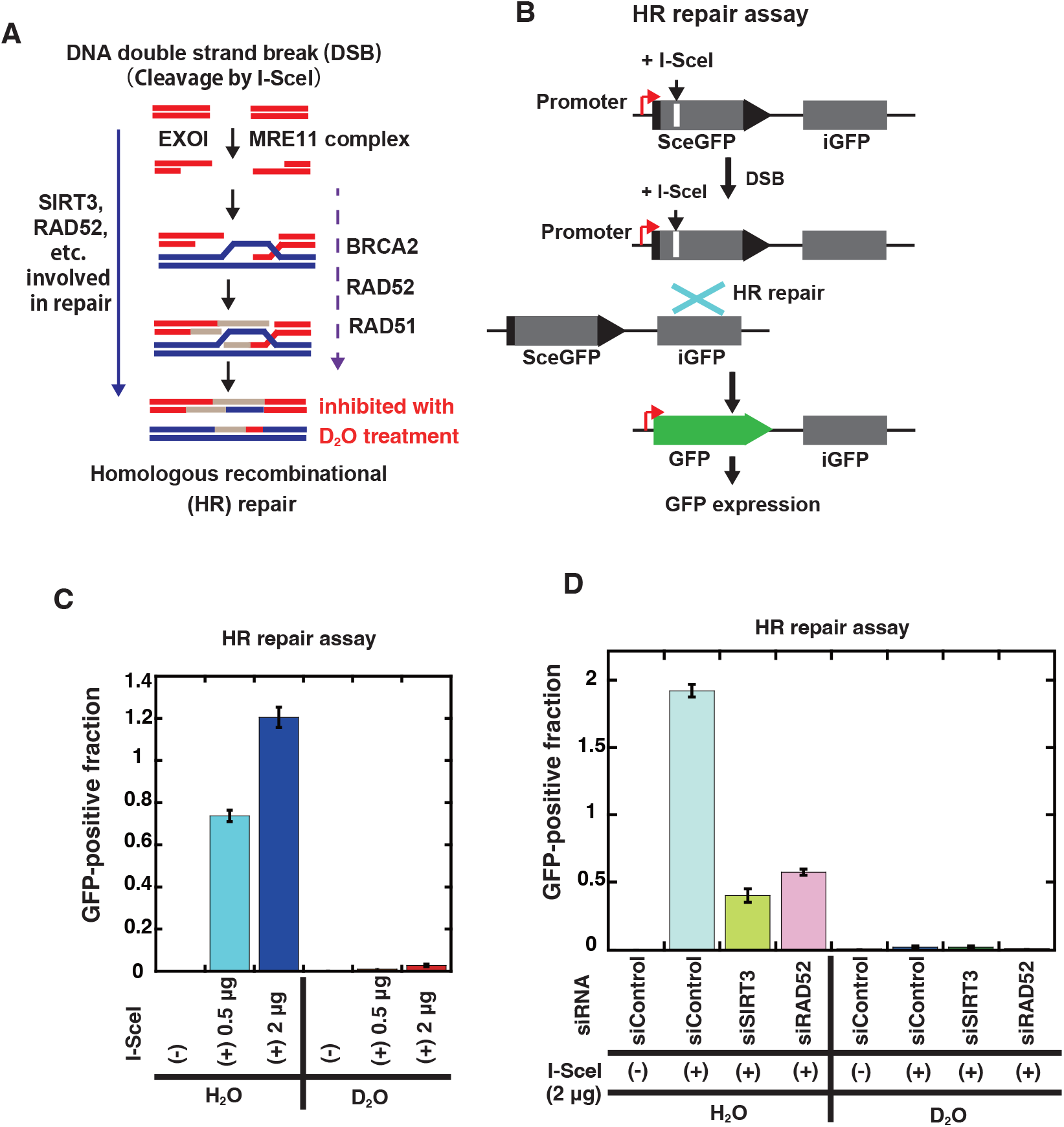
Isotope effects of D_2_O on HR repair in human cells. (**A**) Schematic representation of homologous recombination (HR) repair of an I-SceI-induced DSB site. (**B**) Schematic representation of the I-SceI-induced HR repair reporter assay. HR repair at I-SceI-induced specific DSB sites produces GFP-positive cells. (**C** and **D**) I-SceI-based reporter assays for HR were performed in HeLa pDR-GFP cells cultured in medium containing H_2_O or D_2_O and transfected with the indicated amount of the I-SceI plasmid, as described in the Materials and Methods. The graph shows the mean values and standard errors of the mean from 3 or 6 samples (2 μg of I-SceI in Fig. 4C).

Because SIRT3-mediated RAD52 deacetylation is involved in HR repair, the depletion of RAD52 or SIRT3 decreased the HR repair efficiency (fig. 4D) (*14, 29*). In the *in vitro* experiments, the presence of D_2_O reduced the reaction rate by at most 1/2. Unexpectedly, we found that HR repair was almost completely inhibited in cells cultured in medium containing D_2_O (fig. 4, C and D). HR repair is also required for normal cell growth, and cell death is induced by strongly inhibiting HR repair factors (*30-32*). This might be one of the reasons for our results that cell death was induced by exposure to D_2_O in various types of normal and cancer cells, in a D_2_O concentration-dependent manner, consistent with previous reports (fig. S3, A-D) (*33-35*).

## Discussion

For understanding life phenomena at the quantum level, several books and reviews have described imaginative future possibilities regarding applications of quantum biology to various life phenomena (*1, 36*). The authors suggested that chemical reactions by biomolecules, which make up life, are ultimately quantum level reactions and might therefore be subject to quantum effects. However, some skepticism remains as to whether such microscopic-level effects by quantum mechanics actually have any meaningful impact on macroscopic life systems in living cells or organisms.

In this study, we have presented results suggesting the involvement of quantum tunneling effects in several enzyme-mediated hydrolysis reactions involved in deacetylation, protein cleavage, and DNA cleavage reactions, which are involved in diverse biological phenomena, based on kinetic isotope effects and their temperature dependence. In addition, we have experimentally shown the relevant quantum effects on cells attributed to D_2_O. D_2_O is reportedly cytotoxic (*33-35*), but it was unclear whether this was due to phenomena related to quantum effects, since no experimental reports have shown their relevance. In addition, the involvement of quantum tunneling effects in some enzymes had been shown by *in vitro* experiments (*5-8*), but the biological outcomes attributable to these effects have remained elusive. Thus, this study is the first to experimentally connect the relevant quantum effects *in vitro* and in cells.

Protein labeling using carbon or nitrogen isotopes, which are less prone to quantum tunneling due to their large atomic weights compared to hydrogen, is widely used for proteome analysis by the SILAC method because they have no effects on cells (*37*). The substitution of protein atoms with their isotopes is not considered to significantly affect enzyme activities, because the outer electrons important for enzymatic chemical reactions are the same within the isotope. Therefore, the major effect of the hydrogen isotope on cells is thought to be the kinetic isotope effects caused by the quantum effects.

In our *in vitro* experiments, the rate reduction by D_2_O for the RAD52 deacetylation and I-SceI cleavage reactions was at most 1/2, but the experimental results of HR repair involving these enzyme reactions in cells showed that the recombination frequency was almost completely inhibited by the D_2_O-treatment. Therefore, the quantum effects of D_2_O on cells were much greater than those expected from the *in vitro* experiments. This could be attributed to the fact that cellular reactions are successive reactions by various enzymes, so the final effect might appear as a synergistic reduction in the rates of individual enzymatic reactions due to the kinetic isotope effects. For example, in HR repair reactions, nucleases such as the MRE11 complex and ExoI, which cleave the bonds of DSB-terminal bases (*21-22*), are expected to be affected by the quantum effects in a similar manner to the I-SceI nuclease. In addition to RAD52 deacetylation, various other deacetylation reactions are involved in HR repair, and are also expected to be influenced by the quantum effects. Multiple ATPases, such as RAD51, also perform HR reactions using energy produced by ATP hydrolysis (*21*). Although the quantum tunneling effect in the ATPase-mediated ATP hydrolysis reaction has not been experimentally demonstrated in this study, this reaction may also be affected. Thus, there are many reaction steps in HR repair that could be affected by the kinetic isotope effects. If there are 10 reaction steps subjected to the synergistic isotope effects that reduce the reaction rate by 1/2, then the reaction rate is almost completely impeded by (1/2)^10^ = 1/1,024 = 0.0009765625. For this reason, it can be inferred that the HR repair reaction in the cell was much more strongly inhibited than the extent expected from the *in vitro* kinetic isotope effect.

On Earth, ∼99.972% of the isotopic abundance of hydrogen is H with an atomic mass of 1, and only ∼0.028% of the isotopic abundance is D with an atomic mass of 2 (*38*). Thus, in the “origin of life,” organisms optimized for H were more likely to prevail, since H constitutes most of the hydrogen on Earth. Therefore, in an environment where only H is present as hydrogen, the existence of quantum effects would probably not have an overtly negative effect on the survival of cells. However, in a world where only D exists as hydrogen, which is essentially negligible on the Earth, organisms, including humans, would not be able to survive due to its unexpectedly extensive quantum effects.

## Acknowledgments

We thank Dr. M. Jasin (Memorial Sloan-Kettering Cancer Center, NYC, USA) for the DR-GFP assay materials. We deeply appreciate Dr. S. Hirayama (Emeritus Professor, Kyoto Institute of Technology, Kyoto, Japan) and Dr. H. Kitoh-Nishioka (Kindai University, Osaka, Japan) for helpful instructions and discussions about quantum tunneling and kinetic isotope effects.

## Funding

Provide complete funding information, including grant numbers, complete funding agency names, and recipient’s initials. Each funding source should be listed in a separate paragraph.

Examples

JSPS KAKENHI Grant Numbers 20K12177 (T. Yasuda)

JSPS KAKENHI Grant Numbers 20K08071 (K.T. and T. Yasuda)

JSPS KAKENHI Grant Numbers 22H03743 (W.K. and T. Yasuda)

JSPS KAKENHI Grant Numbers 22K08414 (T.G. and T. Yasuda)

The joint research program of the Biosignal Research Center, Kobe University, Grant Numbers 281004 (T. Yasuda and K.S.)

The joint research program of the Biosignal Research Center, Kobe University, Grant Numbers 201003 (T. Yasuda and K.S.)

## Author contributions

Conceptualization: T. Yasuda

Investigation: T. Yasuda, N.N., T. Yanaka, T.G.

Production of research materials: T. Yasuda, W.K., K.S.

Visualization: T. Yasuda, N.N.

Funding acquisition: T. Yasuda, T.G., W.K., K.S., K.T.

Project administration: T. Yasuda

Writing – original draft: T. Yasuda

Writing – review & editing: W.K., K.S., K.T.

## Data and materials availability

All data are available in the main text or the supplementary materials. The biological materials are available upon request.

## Supplementary Materials

Materials and Methods

Supplementary Text S1 and S2

Figs. S1 to S3

