## Supplementary material for "Cell viability is dominated by quantum effects": Materials and Methods

### Recombinant proteins and reagents

The recombinant human p300 and RAD52 (full length) proteins were purified as described previously (*13-14*). Commercially purchased recombinant SIRT3 protein (#50014, Lot 2003, BPS Bioscience, San Diego, CA, USA) was used as described previously (*14*). The following reagents were purchased: Acetyl Coenzyme A (A2181, Merck & Co., Inc., Whitehouse Station, NJ, USA), $NAD^+$ (N7004, Merck & Co., Inc.), BSA (B9001, New England Biolabs, Ipswich, MA, USA), I-SceI (R0694, New England Biolabs), Caspase 3 active (14-264, Merck & Co., Inc.), $H_2O$ (fig. 1B and C, and #1 in fig. S2; purified with a Milli-Q Synthesis A10 water purification system, Merck & Co., Inc.), $H_2O$ (#2 in fig. S2, and other figures unless otherwise noted; 06442-95, Nacalai Tesque, Inc., Kyoto, Japan), $H_2O$ (#3 in fig. S2; H20MB0501, Merck & Co., Inc.), $D_2O$ (fig. 1B and C, and #1 in fig. S2; 151882, Merck & Co., Inc.), $D_2O$ (#2 in fig. S2, and other figures unless otherwise noted; D214H, Nacalai Tesque, Inc.).

### Immunoblotting and antibodies

Immunoblotting analyses were performed as described previously (*14*). The following antibodies were used: anti-acetyl lysine (#9441, Cell Signaling, Danvers, MA, USA), and anti-RAD52 (sc-8350, Santa Cruz Biotechnology, Dallas, TX, USA).

### *In vitro* acetylation and deacetylation assays

*In vitro* acetylation assays of RAD52 were performed by incubating p300 (0.13 μM) and RAD52 (1.7 μM) proteins in HAT buffer [50 mM Tris-HCl, 1 mM EDTA, 10% glycerol, 1 mM DTT, pH 8.0], made with $H_2O$ (06442-95, Nacalai Tesque, Inc.) or $D_2O$

(D214H, Nacalai Tesque, Inc.), in the presence of Ac-CoA (100 μM) at 30°C for 60 min. Subsequently, a poly dT 68 mer (14.8 μM; synthesized by Operon Biotechnologies, Japan) was added to the reaction mixture, which was further incubated at 30˚C for 10 min to inhibit the acetylation reaction of RAD52, as described (*14*). *In vitro* deacetylation assays of the acetylated RAD52 proteins were then performed by incubating an aliquot of the reaction mixture containing the acetylated RAD52 (0.1 or 0.08 μM) with the indicated amount of recombinant SIRT3 protein in HDAC buffer [25 mM Tris-HCl, 137 mM NaCl, 2.7 mM KCl, 1 mM $MgCl_2$, 0.1 mg/ml BSA, pH 8.0], made with $H_2O$ (06442-95, Nacalai Tesque, Inc.) or $D_2O$ (D214H, Nacalai Tesque, Inc.), in the presence of 500 μM $NAD^+$ at the indicated temperature. The reaction mixtures were subjected to SDS-PAGE, followed by immunoblotting with an anti-acetylated lysine antibody and an anti-RAD52 antibody.

***In vitro* caspase-3-mediated peptide cleavage assay**

*In vitro* caspase-3-mediated DEVD peptide cleavage assays were essentially performed as described previously (*20*), using a Caspase 3/7 Assay Kit (17-367, Merck & Co., Inc.). The active caspase-3 recombinant protein (25 ng) (14-264, Merck & Co., Inc.) was incubated at the indicated temperature with the Ac-DEVD-AMC substrate solution (0.75 μl) (12-541, Merck & Co., Inc.) in 75 μl of 1.5 x Caspase buffer [30 mM PIPES, pH 7.2, 112.5 mM NaCl, 3.75 mM EDTA, 0.15% CHAPS, 11.25% sucrose, 15 mmol DTT], made with $H_2O$ (06442-95, Nacalai Tesque, Inc.) or $D_2O$ (D214H, Nacalai Tesque, Inc.). Following the addition of the Ac-DEVD-AMC substrate, the fluorescence intensity resulting from substrate cleavage was measured with excitation at 355 nm and emission at 460 nm, using Wallac 1420 ARVOsx and ARVO X5 microplate readers (Perkin Elmer,

Waltham, MA, USA).

### *In vitro* I-SceI-mediated DNA cleavage assay

For the *in vitro* I-SceI-mediated DNA cleavage assay, the pGP2 NotI-linearized control plasmid (New England Biolabs), which contains a single I-SceI site, was used as the substrate. The substrate DNA (50 ng) was incubated with the indicated amount of I-SceI enzyme in 10 μl of CutSmart buffer [50 mM potassium acetate, 20 mM Tris-acetate, 10 mM magnesium acetate, 100 μg/ml BSA, pH 7.9], made with $H_2O$ (06442-95, Nacalai Tesque, Inc.) or $D_2O$ (D214H, Nacalai Tesque, Inc.), at the indicated temperature. After the reaction, the samples were analyzed by 0.8% agarose gel electrophoresis with ethidium bromide staining. The gel images were captured using a BioSpectrum Imaging System (UVP, Upland, CA, USA) and an LAS-4000 mini imaging analyzer (Fujifilm, Tokyo, Japan). Quantification was performed using the Multi Gauge software (Fujifilm, Tokyo, Japan).

### Cell culture

HeLa pDR-GFP cells, obtained from Dr. M. Jasin (*27-28*), as well as HEK293, HFL III, and U2OS cells, were cultured in minimum essential medium (MEM) (1030700, Thermo Fisher Scientific, Waltham, MA, USA), supplemented with 10% fetal bovine serum (FBS), 2 mM L-glutamine, and 1% penicillin-streptomycin (PS). For experiments comparing the kinetic isotope effects due to $H_2O$ or $D_2O$ in these cells, the MEM solutions were prepared by dissolving MEM powder (61100, Thermo Fisher Scientific) in $H_2O$ (06442-95, Nacalai Tesque, Inc.) or $D_2O$ (D214H, Nacalai Tesque, Inc.) according to the manufacturer's instructions.

**siRNA treatments**

Stealth Select siRNAs were purchased from Thermo Fisher Scientific. Stealth Select siRNA, HSS118726 (5'-AAUCAGCUCAGCUACAUCCUGCAGG-3'), was used for the siRNA treatment against SIRT3. Stealth Select siRNA, HSS109021 (5'-GGCCAAUGAGAUGUUUGGUUACAAU-3'), was used for the siRNA treatment against RAD52. Stealth RNAi negative control siRNA oligonucleotides (Thermo Fisher Scientific) were used for the negative controls. Lipofectamine RNAiMAX was used for the transfection of Stealth Select siRNAs into cells, according to the transfection protocol (Thermo Fisher Scientific).

**I-SceI-based reporter assay for HR**

Hela pDR-GFP cells ($5x10^5$ cells/well in a 12-well plate) were untreated or transfected with siRNA. At 24 h after the siRNA-transfection, the cells were transfected with 0.5 or 2 μg of the I-SceI expression plasmid, pCMV-NLS-I-SceI (*27-28*), and the culture medium was changed to fresh medium made with $H_2O$ (06442-95, Nacalai Tesque, Inc.) or $D_2O$ (D214H, Nacalai Tesque, Inc.). At 48 h after the plasmid-transfection, the cells were harvested by trypsinization and analyzed with a FACSverse flow cytometer (Becton Dickinson, San Jose, CA, USA). GFP-positive cells were counted by using the FlowJo software (Tomy Digital Biology, Tokyo, Japan).

**Cell survival assays**

To monitor the cell viability by an MTT assay, cells were seeded in 96-well flat-bottom plates (3,000 cells/well) (167008, Thermo Fisher Scientific). After 24 h, the cell culture medium was changed to fresh medium composed of various ratios of $H_2O$ (06442-95, Nacalai Tesque, Inc.) and $D_2O$ (D214H, Nacalai Tesque, Inc.). After 6 days, the cell viability was examined with a ROCHE Cell Proliferation kit I (MTT) (11465007001, Merck & Co., Inc.), according to the manufacturer’s instructions, using an ARVO X5 microplate reader (Perkin Elmer).

**Statistical analysis**

The KaleidaGraph software, version 5.0, was used for statistical analyses. Statistical analyses for multiple comparisons were performed using a one-way ANOVA, as described previously (*29*). Statistical analyses between the data of two groups were performed using an unpaired Student´s t-test, as described previously (*29*).
