## Supplementary Text S1 for "Cell viability is dominated by quantum effects"

According to the textbook by P.W. Atkins (*9*), the quantum tunneling probability of the transmitted wave in fig. S1B is calculated by using the Schrödinger equation, as follows.

The Schrödinger equation is:

$$-(\hbar^2/2m)\, d^2\phi/dx^2 + V\phi = E\phi$$
$$\hbar = h/2\pi$$

where $\phi$ (wave function), $E$ (total energy: motion + potential), $m$ (mass of the particle) $h$ (Planck's constant), and $V$ (potential energy of the particle).

In fig. S1B, the Schrödinger equations for each region are:
Region A (V = 0): $-(\hbar^2/2m)\, d^2\phi/dx^2 = E\phi$
Region B (V > 0): $-(\hbar^2/2m)\, d^2\phi/dx^2 + V\phi = E\phi$
Region C (V = 0): $-(\hbar^2/2m)\, d^2\phi/dx^2 = E\phi$.

The solutions of these equations are:
Region A: $\phi_A(x) = Ae^{ikx} + A'e^{-ikx}$ $\quad k = (2mE/\hbar^2)^{1/2}$
Region B: $\phi_B(x) = Be^{k\prime x} + B'e^{-k\prime x}$ $\quad k' = (2m(V-E)/\hbar^2)^{1/2}$
Region C: $\phi_c(x) = Ce^{ikx} + C'e^{-ikx}$ $\quad k = (2mE/\hbar^2)^{1/2}$.

Since there are no particles moving in the − x direction on the right side,
$C' = 0$.
The probability of the quantum tunneling effect, $P$, where a particle passes through a barrier, is

$$P = |C|^2/|A|^2$$

Since $\phi$ and its derivative $d\phi/dx$ are continuous on the left (x = 0) and right (x = L) sides,
$(\phi_A(0) = \phi_B(0);\ (\phi_A/dx)_{x=0} = (\phi_B/dx)_{x=0}$
$\phi_B(L) = \phi_C(L);\ (\phi_B/dx)_{x=L} = (\phi_C/dx)_{x=L}$.
Therefore,
$A + A' = B + B'$; $ikA - ikA' = Bk' - B'k'$
$Be^{k\prime L} + B'e^{-k\prime L} = Ce^{ikL}$; $k'Be^{k'L} - k'B'e^{-k\prime L} = ikCe^{ikL}$

$$P = 1/(1+G)$$

$$G = \frac{[\exp\left[\{2m(V-E)/\hbar^2\}^{1/2}L\right] - \exp\left[-\{2m(V-E)/\hbar^2\}^{1/2}L\right]]^2}{4\,(E/V)\{1-(E/V)\}}$$

If E < V; e.g., E/V = 0.5, then 4 $(E/V)\{1-(E/V)\} = 1$.

Comparing $^{1}$H and $^{2}$H, we get

$$\exp\left[\{2 \times 2 \times (V-E)/\hbar^2\}^{1/2}L\right] > \exp\left[\{2 \times 1 \times (V-E)/\hbar^2\}^{1/2}L\right]$$

$$\exp\left[-\{2 \times 2 \times (V-E)/\hbar^2\}^{1/2}L\right] < \exp\left[-\{2 \times 1 \times (V-E)/\hbar^2\}^{1/2}L\right].$$

Therefore,

$$G_{^1H} < G_{^2H}$$

and

$$P_{^1H} > P_{^2H}$$

Thus, $^{1}$H has a higher probability of quantum tunneling than $^{2}$H.
