## Supplementary Text S2 for "Cell viability is dominated by quantum effects"

The vibrational potential energy of a chemical bond involved in a reaction is described as follows.

$$\nu = \frac{1}{2\pi}\sqrt{\frac{\kappa}{\mu}}$$

(where $\nu$ is the vibration frequency of the chemical bond between two atoms, and $\kappa$ is the force constant of the chemical bond),

$\mu = (m \times M) \div (m + M)$

(where $\mu$ is the reduced mass, and m and M are the masses of the two atoms), and

$$E_n = \left(n + \frac{1}{2}\right) h\nu$$

(where $E_n$ is vibrational potential energies according to the quantum mechanics, n = 0, 1, 2, 3, etc., and h is the Planck constant).

Therefore, the lowest vibrational potential energy is described as below.

$$E_0 = \frac{1}{2} h\nu = \frac{h}{4\pi}\sqrt{\frac{\kappa}{\mu}}$$

(where $E_0$ is the vibrational zero-point energy).

Accordingly, the zero-point energy ($E_0$) is decreased when the reduced mass ($\mu$) is increased, and the reduced mass is increased when the mass of one of the atoms involved in the chemical reaction is increased. Assuming that one of the atoms is hydrogen (H), the $E_0$ value is changed at both the GS and TS when the hydrogen is substituted with its heavier isotope deuterium (D) (Figure 1c). Therefore, the activation energy ($\Delta E$) is increased by the substitution of H with D, thereby decreasing the reaction rate by the substitution.
