## Supplementary material for "Cell viability is dominated by quantum effects": FigS1

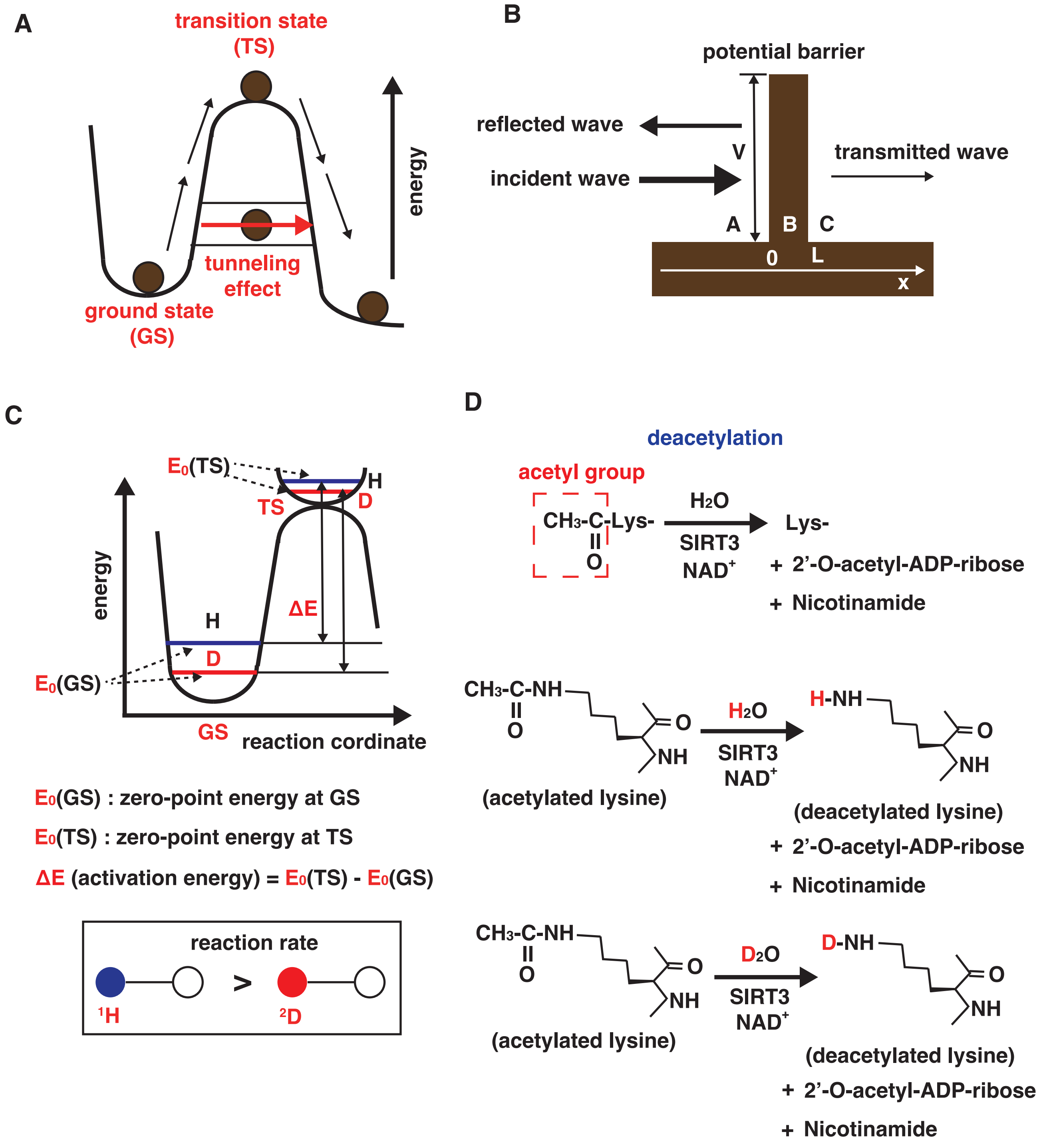


**Fig. S1. Schematic representations of the concept of this research.** (**A**) Conceptual image of quantum tunneling, in which the chemical reaction of the reactant (brown circle) at the ground state can proceed without reaching the transition state by penetrating through the energy barrier. (**B**) Conceptual diagram of quantum tunneling with a 1D box barrier model (*9*). The height of the potential barrier (V), the thickness of the potential barrier (L), and the incident wave, reflected wave, and transmitted wave are shown. (**C**) Difference in zero-point vibrational energies between hydrogen (H) and deuterium (D) to explain the kinetic isotope effect (*6,7,9*). (**D**) Chemical reactions of SIRT3-mediated deacetylation of an acetylated lysine residue in the presence of $H_2O$ or $D_2O$.
