## Supplementary material for "Cell viability is dominated by quantum effects": FigS2

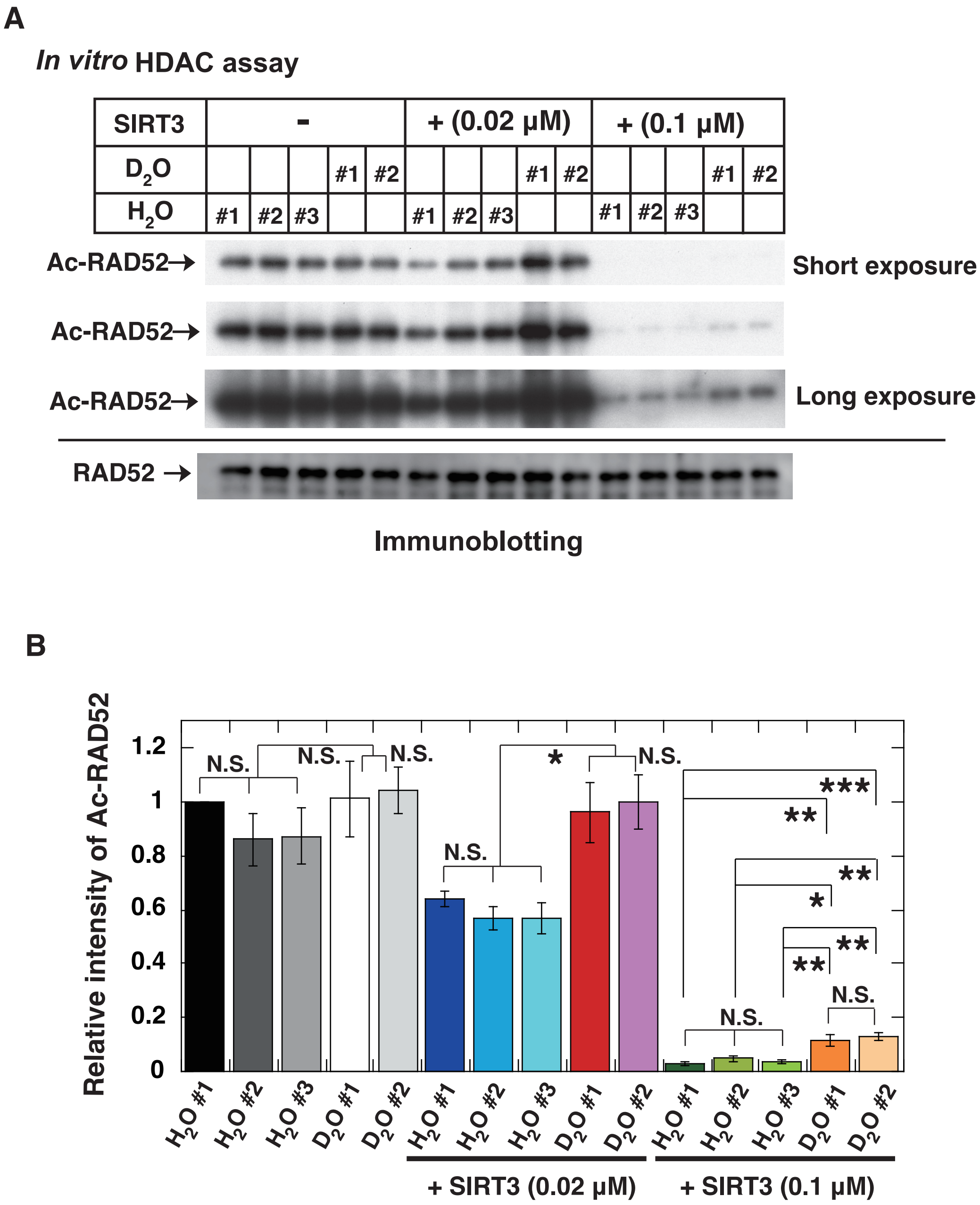


**Fig. S2. Kinetic isotope effect of SIRT3-mediated deacetylation of RAD52 in the presence of $H_2O$ and $D_2O$ from different sources.** (**A**) In vitro acetylation assays were performed as described in Fig. 1, in the presence of $H_2O$ or $D_2O$ purchased from different sources. $H_2O$ #1, $H_2O$ #2, $H_2O$ #3, $D_2O$ #1, and $D_2O$ #2 were used, as described in the Materials and Methods. After the addition of a poly dT 68 mer, an aliquot of the reaction mixture containing the RAD52 protein (final concentration 0.08 μM) was incubated with the indicated amount of SIRT3 in HDAC buffer, prepared with $H_2O$ or $D_2O$ from different manufacturers, at 30˚C for 60 min. The reaction mixtures were subjected to SDS-PAGE, followed by immunoblotting with an anti-acetylated lysine antibody (Ac-RAD52) and an anti-RAD52 antibody (bottom, RAD52). The long and short exposures are also presented for the immunoblotting with an anti-acetylated lysine antibody. (**B**) The relative band intensities of acetylated RAD52 normalized to those of the RAD52 bands are shown in the graph. The mean values and standard errors of the mean from 3 independent experiments were plotted. For each SIRT3 concentration, the samples connected by lines were compared (*P <0.05, **P <0.01, **P <0.001 and N.S., not significant by one-way ANOVA with Dunnet's post hoc test with the KaleidaGraph software).
