## Supplementary material for "Cell viability is dominated by quantum effects": FigS3

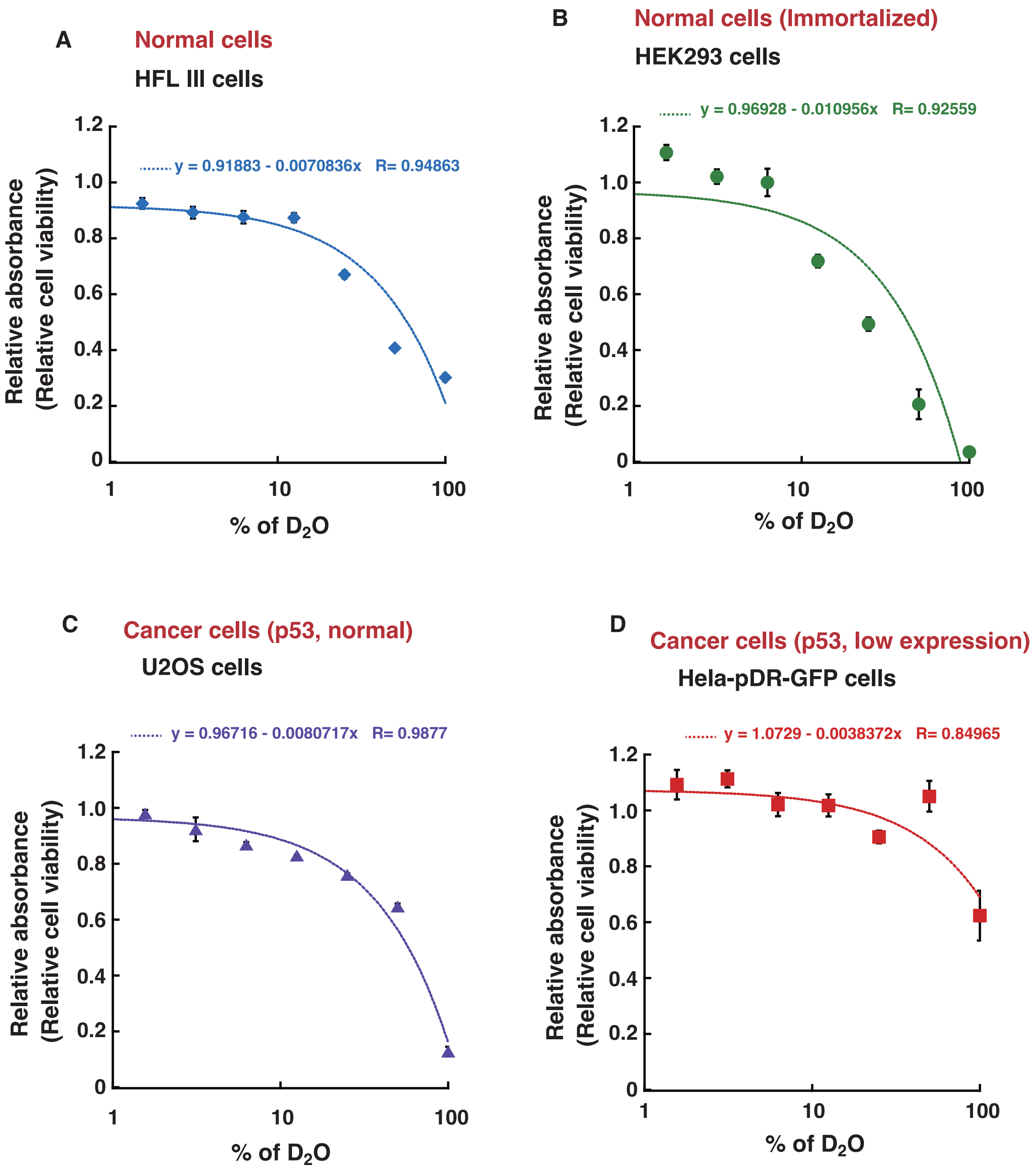


**Fig. S3. Cytotoxicity of $D_2O$ in human cells.** (**A** to **D**) HFL III (A), HEK293 (B), U2OS (C), and HeLa pDR-GFP (D) cells were cultured in media containing various concentrations of $D_2O$ for 6 days. Cell survival was examined by an MTT assay, as described in the Materials and Methods. The graph shows the mean values and standard errors of the mean from 3 samples, with linear curve fitting.
